# A functional anatomical shift from the lateral frontal pole to dorsolateral prefrontal cortex in emotion action control underpins elevated levels of anxiety – partial replication and generalization of Bramson et al., 2023

**DOI:** 10.1101/2025.03.09.642129

**Authors:** Qian Zhuang, Shuxia Yao, Lei Xu, Shuaiyu Chen, Jialin Li, Xiaoxiao Zheng, Meina Fu, Keith M. Kendrick, Benjamin Becker

**Affiliations:** Center for Cognition and Brain Disorders, The Affiliated Hospital of Hangzhou Normal University, Hangzhou, Zhejiang Province, China; The Clinical Hospital of Chengdu Brain Science Institute, MOE Key Laboratory for Neuroinformation, Center for Information in Medicine, University of Electronic Science and Technology of China, Chengdu, China; Institute of Brain and Psychological Sciences, Sichuan Normal University, Chengdu, China; Brain Cognition and Brain Disease Institute (BCBDI), Shenzhen Institute of Advanced Technology, Chinese Academy of Sciences, Shenzhen, China; Institute of Science and Technology for Brain-Inspired Intelligence, Fudan University, Shanghai, China; State Key Laboratory of Brain and Cognitive Sciences, The University of Hong Kong, Hong Kong, China; Department of Psychology, The University of Hong Kong, Hong Kong, China

**Keywords:** Emotion, inhibition, anxiety, frontal pole, DLPFC, Go/NoGo task

## Abstract

Flexible control over emotional behavior represents a promising target for novel interventions for mental disorders. Accumulating evidence has indicated a key role of the lateral frontal pole (FPl) and its connections with other cortical and subcortical systems in emotional action regulation. A recent study from Bramson et al., (2023) employed a multi-modal neuroimaging approach to demonstrate a functional-anatomical shift from FPI to dorsolateral prefrontal cortex (DLPFC) in a sample of anxious individuals during emotional action control. While these findings might represent a venue for interventions in anxiety disorders, conventional neuroimaging strategies are often limited with respect to generalizability and reproducibility. Against this background we capitalized previous large-scale fMRI data in n = 250 participants using an affective linguistic Go/NoGo paradigm to examine the robustness of the reported associations with trait social anxiety across samples, cultures and paradigms. Additionally, context-dependent functional connectivity patterns were explored to examine action control in different emotional contexts. In line with previous study, we found no difference between high- and non-anxious group on the behavioral congruency-effect. The neural results showed that non-social anxious group engaged the left FPl while the high-social anxious group specifically recruited the DLPFC, however in the absence of significant between-group differences. Importantly, the level of trait social anxiety was significantly positively related with DLPFC activity and negatively with left FPl activation across groups. Furthermore, context-dependent functional connectivity analyses revealed a negative context-specific neural shift from the sgACC-FPl to sgACC-DLPFC specifically in the high anxiety group. Together, the present study employed a different task paradigm, population and analytic methods, partially replicated the findings described by Bramson et al., (2023) and additionally determined context-specific changes in the communication with the sgACC in high anxiety. The findings provide further evidence for target-based interventions of persistent emotional control deficits in anxiety disorders.

---

Flexible control over emotional behavior is vital for mental health and represents a promising target for novel interventions for mental disorders (e.g. Etkin et al., 2015; Feng et al., 2018). Accumulating evidence from task-based neuroimaging studies indicates a key role of the lateral frontal pole (FPl) and its connections with other cortical and subcortical systems in regulating emotional action tendencies (Bramson et al., 2023; Fonzo et al., 2017; Tyborowska et al., 2016, 2024). A recent study from Bramson et al., (2023) employed a multi-modal neuroimaging approach to demonstrate a functional-anatomical shift in a sample of anxious individuals, such that high-anxiety individuals engaged the dorsolateral prefrontal cortex (DLPFC), while non-anxious individuals engaged the FPI during control of emotional action tendencies. While these findings might represent a venue for interventions targeting persistent emotional control deficits in anxiety disorders (Meijer et al., 2023), conventional neuroimaging strategies are often limited with respect to generalizability and reproducibility (e.g. Gan et al., 2024; Poldrack et al., 2017; Zhou et al., 2022). Against this background we here capitalized on data from a previous large-scale fMRI study from our team (Zhuang et al., 2021, 2023) in n = 250 healthy individuals using a motor control task (affective linguistic Go/NoGo paradigm, Fig.1A) to examine the robustness of the reported associations with social anxiety across samples, cultures and paradigms (details see Supplementary Information (SI)). The large dataset allowed us to split the sample into high- and low-social anxiety groups and the paradigm allowed us to model emotional action control as the capability to override the automatic action tendencies evoked by emotional words in affect-incongruent conditions (Happy NoGo (HNG) and Fearful Go (FG) trials) as compared to congruent conditions (Happy Go(HG) and Fearful NoGo(FNG) trials) during the emotion and inhibitory control interactions according to a well-established Pavlovian bias framework (Guitart-Masip et al., 2012, 2014). Additionally, we explored emotional context-dependent (contrasts: FG>FNG; HNG>HG) functional connectivity patterns to examine action control in different emotional contexts.

**Fig. 1.**
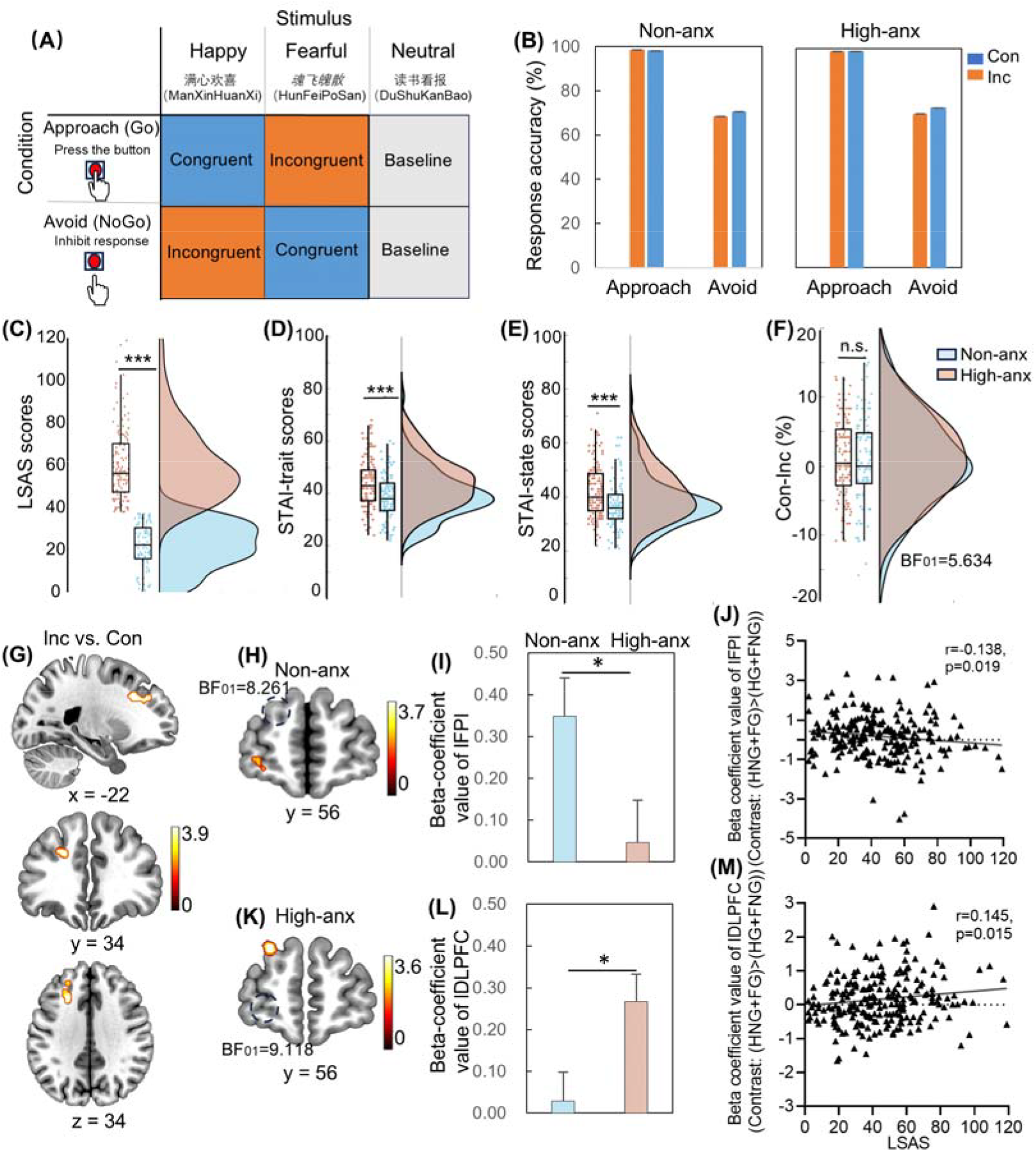
Behavioral and neural-congruence effect during emotional behavioral control with the affective Go/NoGo paradigm. (A) Congruent-incongruent effect as modelling by the affective Go/NoGo task used in the current study to delineate the interaction between emotion and behavioral factors. (B) Mixed ANOVA analysis on response accuracy with congruence (incongruent/congruent) *action (approach/avoid)*group (high-anxious/non-anxious) as variables showed a main effect of congruence (F_(1,225)_=6.174, p=0.014) and action (F_(1,225)_=692.071, p<0.001). Neither the interaction effect between congruence*action*group (F_(1,225)_=0.001, p=0.970) nor the interaction between congruence*group (F_(1,225)_=0.415, p=0.520) was significant with (F) the Bayesian t-test showing a moderate evidence for the absence of group difference on behavioral congruence effect (BF_01_=5.634). (C) Group differences on LSAS, TAI (D) and SAI (E) with LSAS 38 as cutoff score. (G) Neural-congruence effect across groups showed a significant activation in left DLPFC after small volume correction. (H&K) Examination of the anxiety groups with one sample tests (based on contrast: (HNG+ FG) > (HG+ FNG)) confirmed that the non-social anxious group mainly engaged the left FPl (small volume correction (SVC), Z=3.54, p_FWE_=0.039, voxels=12, x/y/z: -30, 60, -3), while the high-social anxious group specifically recruited the DLPFC (SVC, Z=3.47, p_FWE_=0.048, voxels=13, x/y/z: -27, 42, 36; threshold: SVC in combination with peak-level FWE corrected at p < 0.05). (I&L) Parameter estimates (contrast: (HNG+ FG) > (HG+ FNG)) extracted by the activated region confirmed group differences on the BOLD signal of left FPl and DLPFC with stronger recruitment of left FPl and DLPFC in the non-social anxious group and high-social anxious group, respectively. (J&M) Brain BOLD signal (contrast: (HNG+ FG) > (HG+ FNG)) and trait anxiety association analyses showed a significant negative association between left FPl activation and LSAS scores (r=0.138, p=0.019) and a positive association between left DLPFC and LSAS scores (r=0.145, p=0.015) across groups (following a Bonferroni correction p<0.05/4=0.0125, trend-to-significant) but not for STAI scores (all ps>0.226). Note: Con: congruent; DLPFC, dorsolateral prefrontal cortex; FPl, lateral frontal pole; High-anx: high-social anxious group; Inc: incongruent; non-anx: non-social anxious group. * means p<0.05, *** means p<0.001, n.s means non-significant.

After quality control n=227 participants were divided into high- (n=127) and non-social anxious group (n=100) using the Liebowitz Social Anxiety Scale (LSAS, Fig.1C) with 38 as cutoff score (based on He & Zhang, 2004, reporting both sensitivity and specificity>80% in a Chinese population). Groups were further validated with STAI-trait (p<0.001, Fig.1D) and state anxiety differences (p<0.001, Fig.1E, Spielberger, 1983).

Mixed ANOVA analysis on behavioral response accuracy showed no significant interaction effect (F_(1,225)_=0.001, p=0.970) but a main effect of action (F_(1,225)_=692.071, p<0.001) and congruence (F_(1,225)_=6.174, p=0.014, Fig. 1B, see SI), which was in line with the findings from Bramson et al., (2023). Additionally, in line with Bramson et al., (2023), the difference between high- and non-social anxious group on the behavioral congruency-effect was not significant (F_(1,225)_=0.415, p=0.520) with a Bayesian t-test showing moderate null hypothesis evidence (BF_01_=5.634, Fig.1F).

The neural results found a significant neural congruency-effect in the left DLPFC (SVC, Z=3.79, p_FWE_=0.015, voxels=40, x/y/z: -21, 24, 33, Fig.1G) across groups. In line with Bramson et al., (2023), examination of the anxiety groups confirmed that the non-social anxious group engaged the left FPl (small volume correction (SVC), Z=3.54, p_FWE_=0.039, voxels=12, x/y/z: -30, 60, -3. Fig.1H &I), while the high-social anxious group specifically recruited the DLPFC (SVC, Z=3.47, p_FWE_=0.048, voxels=13, x/y/z: -27, 42, 36. Fig.1K & L). While no significant between-group differences were found, further Bayesian t-tests resulted in moderate evidence for the absence of left FPl activation in the high-social anxious group (BF_01_=9.118) as well as the left DLPFC in the non-social anxious group (BF_01_=8.261) during emotional behavior control. In addition, in the entire sample the level of trait social anxiety was significantly positively related with DLPFC activity (r=0.145, p=0.015, Fig.1M) and negatively with left FPl activation (r=0.138, p=0.019, Fig.1J) across groups (following a Bonferroni correction p<0.05/4=0.0125, trend-to-significant) but not with STAI scores (all ps>0.226) as reported by Bramson et al., (2023).

Based on previous studies showing the contribution of intrinsic and emotion-specific network level changes to elevated anxiety (Chen et al., 2023; Xu et al., 2021) and the key role of interactions between the subgenual anterior cingulate cortex (sgACC) with medial and lateral PFC regions in emotion-related cognitive action control (Lapate et al., 2022) we examined whether the network level interaction of these regions varies as a function of anxiety levels. Examining the corresponding contrast in the non- and high-social anxious group separately and focusing on the sgACC-FPl and sgACC-DLPFC circuitry (Methods see SI), we found a significant positive connectivity between left sgACC (seed defined from the contrast: FG>FNG) and bilateral DLPFC as well as a negative connectivity with bilateral FPl for the high-social anxious group specific to the negative context (Fig. 2A, results in the positive context see Fig. 2B) after SVC (initial thresholding: p<0.001, uncorrected; SVC: left DLPFC, Z=5.22, p_FWE_ <0.001, voxels=96, x/y/z: -21, 39, 39, Fig. 2C; right DLPFC, Z=3.79, p_FWE_ =0.031, voxels=28, x/y/z: 18, 30, 39, Fig. 2E; left FPl, Z=4.16, p_FWE_ =0.008, voxels=53, x/y/z: -33, 51, 12, Fig. 2D; right FPl, Z=4.12, p_FWE_ =0.016, voxels=37, x/y/z: 33, 51, 15, Fig. 2F), potentially reflecting network level markers for increased social anxiety. Across both groups the level of social anxiety was positively associated with sgACC-bilateral DLPFC connectivity strength (sgACC-left DLPFC: r=0.193, p=0.002, Fig.2C; sgACC-right DLPFC: r=0.177, p=0.004, Fig.2E) but not with sgACC-FPl connectivity after the multiple comparison corrections (sgACC-left FPl: r=0.055, p=0.206, Fig.2D; sgACC-right FPl: r=0.119, p=0.037, Fig.2F). In addition, no significant correlations between STAI scores and the sgACC-FPI and sgACC-DLPFC connectivity were found (ps≥ 0.016). For the sgACC seed from the contrast: HNG>HG, there was a significant positive connectivity with bilateral DLPFC, but not FPl after SVC, specifically for the high-social anxious group in the negative context (see SI). Together these findings may reflect a negative context-specific neural shift in terms of functional connectivity from the sgACC-FPl to sgACC-DLPFC in high social anxiety during emotional action control. In contrast to Bramson et al., (2023), a direct comparison in the present data did not reveal significant neural differences between the groups, such that differences in the neural congruency effect were mainly reflected in within-group contrasts. However, correlation analyses indicate supporting associations with the levels of social anxiety in the entire sample indicating a partial replication (possible due to the higher sensitivity of dimensional as compared to categorical analyses, see e.g. Chen et al., 2023). Additional analyses revealed convergent network level markers which may reflect that context-specific changes in the communication with the sgACC may underpin deficient emotion regulation in high anxiety.

**Fig. 2.**
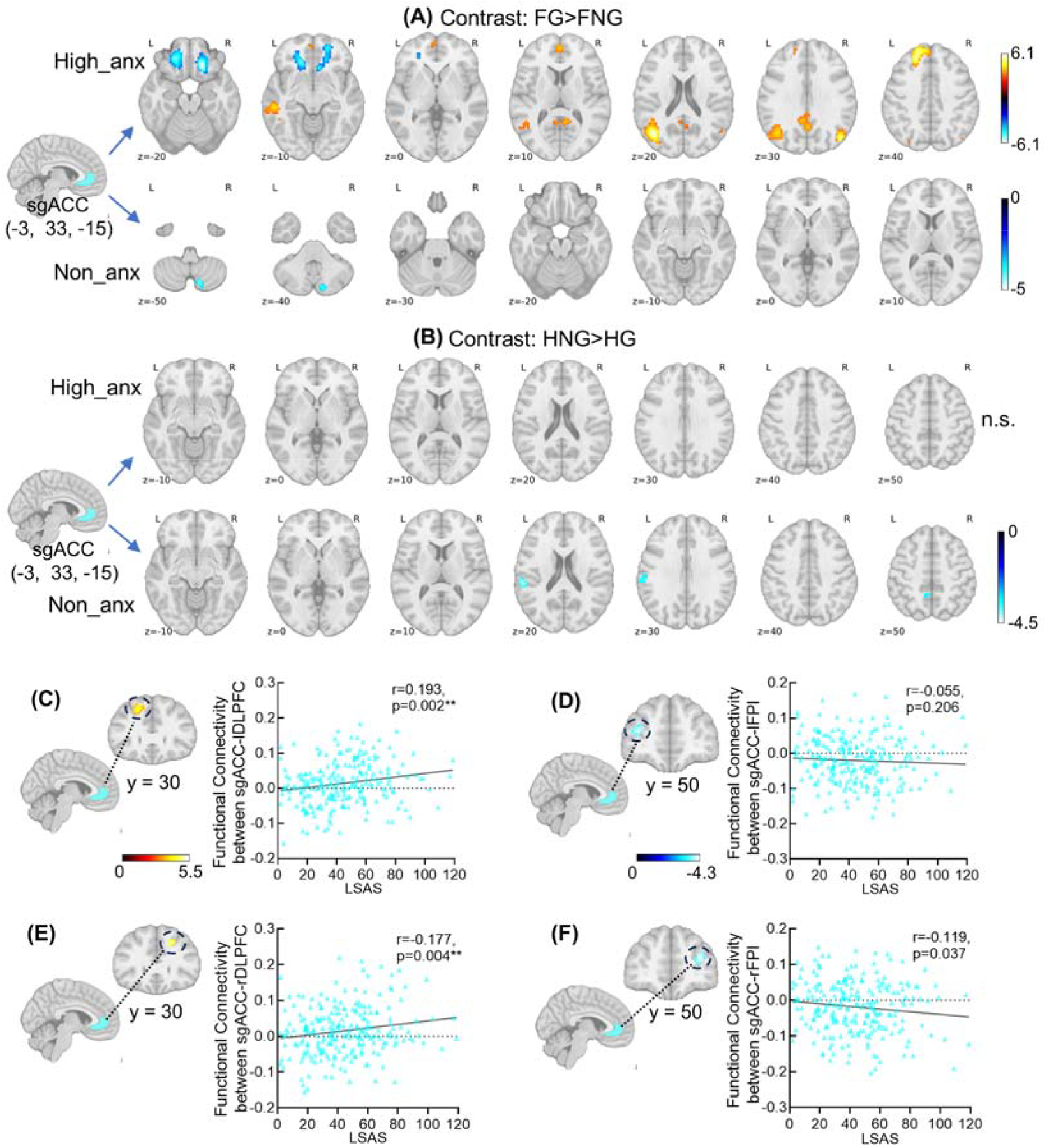
Functional connectivity results in positive and negative emotional context. (A)The results revealed a significant positive connectivity between left sgACC (seed defined by the contrast: FG>FNG) and bilateral DLPFC as well as a significant negative connectivity with bilateral FPl for the high-anxious group rather than the non-anxious group especially in the negative context after small volume correction with corresponding masks centered into the peak coordinates reported by Bramson et al., (2023). (B) No significant connectivity between sgACC (seed defined by the contrast: FG>FNG) and lateral frontal areas were found in the positive context neither for high-anxious nor non-anxious group. The brain and scales association analyses across groups showed significant positive correlations between LSAS scores with sgACC-bilateral DLPFC connectivity strength in the negative context (sgACC-left DLPFC: r=0.193, p=0.002, Fig.2C; sgACC-right DLPFC: r=0.177, p=0.004, Fig.2E) but not with the sgACC-bilateral FPI connectivity strength after the correction for multiple comparisons (sgACC-left FPl: r=0.055, p=0.206, Fig.2D; sgACC-right FPl: r=0.119, p=0.037, Fig.2F). Note: DLPFC, dorsolateral prefrontal cortex; FPl, lateral frontal pole; High-anx: high-social anxious group; non-anx: non-social anxious group; sgACC, subgenual anterior cingulate cortex. ** means p<0.01.

## Supporting information

Supplemental material for Table and Figures

## Conflicts of interests

The authors declare no competing interests.

## Acknowledgements

National Natural Science Foundation of China (grant numbers 82271583 – BB, 31530032 – KMK, 32200904 – Q Z), Key Technological Projects of Guangdong Province (grant number 2018B030335001 – KMK) and Medical and Health Technology Project of Zhejiang Provincial Health Commission (2023RC236 – Q Z).

## Notes

### Competing Interest Statement

The authors have declared no competing interest.

