## Supplemental material for Table and Figures for "A functional anatomical shift from the lateral frontal pole to dorsolateral prefrontal cortex in emotion action control underpins elevated levels of anxiety – partial replication and generalization of Bramson et al., 2023"

Zhuang., et al.,

Contact:

1. Method

1.1 Participants and task setting

N = 250 healthy, right-handed Chinese students were enrolled in the current study after given written informed consent and performed an affective Go/NoGo task with concurrent fMRI (details see Zhuang et al., 2021, 2023). Positive (happy), negative (fearful) and neutral words (word length=4, words frequency matched) were used as task stimuli. For example, while the words included Huan Tian Xi Di (欢天喜地) and Xi Chu Wang Wai (喜出望外) were used on the positive condition, Xin Jing Rou Tiao (心惊肉跳) and Ti Xin Diao Dan (提心吊胆) were used on the negative condition and Che Lai Che Wang (车来车往) and Xin Wen Bo Bao (新闻播报) were used on the neutral condition respectively. Participants were instructed to respond according to the font of the words regardless of the meaning, such as, pressing the button when the words were presented as normal font (i.e. go trials, approach condition), while withdrawing the response when the words were occasionally in italic font (i.e. nogo trials, avoid condition). Among this, while happy Go trials and fearful NoGo trials represented approaching- and avoiding-congruent condition respectively, fearful Go and happy NoGo trials represented approaching- and avoiding-incongruent trials.

During the data quality control, n=23 participants were excluded due to technical issues or excessive head motion during data acquisition. In detail, data from 11 subjects were lost due to technical issues during fMRI (n=5) and behavioral data collection (n=6) and data from another 12 subjects were further excluded due to excessive head motion (>2.5mm or 2.5 degrees, n=12). Thus, there are n = 227 subjects were included in the final behavioral and fMRI data analyses (112 males; age: mean ± SD=21.62 ± 2.32 years) in which n=9 subjects with only first run data. All participants were divided into high-anxious (n=127, 64 female, age: mean+SEM=21.73+0.20) and non-anxious (n=100, 51 female, age: mean+SEM=21.48+0.24) group according to the validated Liebowitz Social Anxiety Scale (LSAS, Mennin et al., 2002; subscale fear: Cronbach’a =0.937; avoid: Cronbach’a=0.917) with 38 as the cutoff score, which has been showed to be with a good sensitivity (85.5%) and specificity (81.3%) in screening for social anxiety disorder in Chinese population (He & Zhang, 2004). There was a significant group difference on the independent measurement of both trait (high-anx: mean+SEM=44.49+0.83; non-anx: mean+SEM=39.05+0.93, t_(225)_=4.37, p<0.001) and state anxiety (high-anx: mean+SEM=41.94+0.86; non-anx: mean+SEM=37.55+0.93, t_(225)_=3.44, p<0.001) scores assessed by State Trait Anxiety Inventory (STAI, Spielberger, 1983). The study was approved by the local ethics committee and in accordance with the latest revision of the Declaration of Helsinki.

2. Data acquisition and analyzing

2.1 Behavioral data analysis

The behavioral data were analyzed with SPSS Version 25.0 (Armonk, NY: IBM Corp.). For response accuracy, the mixed three-way ANOVA analysis with congruence (incongruent/congruent) * action(approach/avoid) * group(high-anxious/

non-anxious) as variables was conducted to examine the group difference on behavioral performance during action control. In addition, for response reaction time on the correct Go trials (e.g. approach trials) the two-way mixed ANOVA analysis with congruence(incongruent/congruent) * group(high-anxious/non-anxious) as variables was conducted. Furthermore, the group differences on the behavioral congruency-effect were conducted with independent samples t-tests. In details, the behavioral congruency-effect was calculated as the response accuracy in the congruent condition (approaching happy and avoiding fearful stimuli) minus the incongruent condition (approaching fearful and avoiding happy stimuli). To further test the null hypotheses that there were no group differences on the behavioral and neural congruency-effect, the Bayesian independent t-tests were conducted using JASP (https://jasp-stats.org).

2.2 MRI Data Acquisition

Neuroimaging data were collected using a 3T GE Discovery MR750 system (General Electric Medical System, Milwaukee, WI). A total of 488 volumes of T2*-weighted echo planar images were acquired (acquisition parameters: repetition time, 2000ms; echo time, 30 ms; slices, 39; slice-thickness, 3.4mm; gap, 0.6mm; field of view, 240 × 240 mm^2^; matrix size, 64 × 64; flip angle, 90°). To improve normalization of the functional images and identify individuals with apparent brain pathologies, high-resolution whole brain T1-weighted images were obtained using a 3D spoiled gradient echo pulse sequence (acquisition parameters: repetition time, 6ms; echo time, minimum; flip angle, 9°; field of view = 256 × 256mm; acquisition matrix, 256 × 256; thickness, 1mm; 156 slices). OptoActive MRI headphones (http://www.optoacoustics.com/) were used to reduce acoustic noise during MRI data acquisition.

2.3 MRI Data analysis

2.3.1 fMRI Data Preprocessing

Functional MRI data were preprocessed using SPM12 software (Wellcome Trust Center of Neuroimaging, University College London, London, United Kingdom). The first 10 volumes for each run were deleted to allow magnet steady data. The remaining functional images were processed using the following standard preprocessing procedures: (1) slice-timing and head motion correction, (2) spatial normalization to Montreal Neurological Institute (MNI) standard space (by means of co-registration to the T1-weighted structural images and the application of the transformation matrices obtained from the segmentation of the structural images), (3) resampling with a 3 x 3 x 3 mm resolution, and (4) spatial smoothing using a 8mm full-width at half-maximum (FWHM) Gaussian kernel.

2.3.2 BOLD level analysis

The general linear model was conducted for the first level analyses using an event-related analysis approach. Six condition-specific regressors (Happy Go (HG, i.e. approaching-congruent), Happy NoGo (HNG, i.e. avoiding-incongruent), Fearful Go (FG, i.e. approaching-incongruent), Fearful NoGo (FNG, i.e. avoiding-congruent), Neutral Go (NeuG), Neutral NoGo (NeuNG)) were modelled on the first level and convolved with the standard hemodynamic response function (HRF). To further control for head movement-related artifacts the six head-motion parameters were included as nuisance regressors. Then, contrast images of interest (neural congruency-effect: (HNG+ FG) > (HG+ FNG); emotional context-specific contrasts: HNG > HG; FG > FNG) were created on the individual level. In the second level model analysis, to examine the general neural-congruent effect across groups as well as group differences on it, one sample t tests and two sample t tests on (HNG+FG) > (HG+FNG) contrasts were computed respectively. In addition, to examine the neural mechanism underlying action control in the positive and negative context separately as well as group differences on it, one sample t tests and two sample t tests on emotion-specific contrasts (HNG > HG; FG > FNG) were computed respectively.

2.3.3 Functional connectivity analysis

To examine whether the connectivity patterns between sgACC and lateral PFC subregions, i.e. DLPFC and FPl, would be shifted as the function of different anxiety level especially in the negative context, the functional connectivity analyses were conducted in each group using generalized psychophysiological interaction analyses (gPPI) approach (McLaren et al., 2012) implemented in CONN toolbox with sgACC as seeds in the positive and negative context separately. The seeds were defined as 6mm spheres centered at the peak coordinates of sgACC activation mapping (sgACC mask from Human brainnetome atlas (Fan et al., 2016)) from the negative (contrast: FG>FNG) and positive condition (contrast: HNG>HG) across groups (details of sgACC activation in different emotional contrasts see Table. S3). The definition for regressors during the gPPI analyses included psychophysiological interactions for HG condition (i.e. approaching-congruent), HNG condition (i.e. avoiding-incongruent), FG (i.e. approaching-incongruent), FNG condition (i.e. avoiding-congruent), NeuG and NeuNG condition, task regressors for the above six conditions respectively, seed region timecourse (i.e. left or right sgACC), and a constant, which allows the testing of any combination of all conditions (McLaren et al., 2012).

2.3.4 Thresholding and regions of interest analyses

The main aim of the current study using the affective Go/NoGo task with big-sample size was to validate the shifted role from FPl to DLPFC in controlling of emotional behaviors for high-anxious individuals and secondly to examine their related functional connectivity patterns in different emotional contexts. To allow a more sensitive examination of the neural-congruence effect on FPl and DLPFC, ROI analyses were conducted with small volume correction (SVC, initial thresholding: p<0.001, uncorrected; peak-level, p_FWE_ < 0.05) using the 15mm spheres as masks which is centered at the coordinates reported by Bramson et al., (2023). In addition, the whole brain cluster-level correction was employed on the BOLD level and functional connectivity analyses (initial thresholding: p<0.001, cluster-level, p_FWE_<0.05).

2.4 Brain and trait association analyses

Brain and scales association analyses including LSAS and total STAI scores were performed with age and gender as covariates, in which one-tailed significance tests (p<0.05) were set based on previous findings showing less FPl instead of more DLPFC recruitment in emotional action control as the increased anxiety level (Bramson et al., 2023). For the brain activation of the left FPl and DLPFC, parameter estimates were extracted as mean values of voxels within the activated regions in each condition using the scripts mainly encompassed the spm_select and spm_summarise function implemented in Matlab (R2018a). Additionally, for the functional connectivity between sgACC and FPl and DLPFC, parameter estimates were extracted with the same method. The correction for multiple comparisons for the correlation analyses in respect of brain activity (p<0.05/4=0.0125) and functional connectivity (p<0.05/8=0.006) was employed.

3. Results

3.1 Behavioral results

For the response accuracy, the mixed ANOVA analysis with congruence (incongruent/congruent) *action (approach/avoid)*group (high-anxious/non-anxious) as variables showed a main effect of congruence (F_(1,225)_=6.174, p=0.014) and action (F_(1,225)_=692.071, p<0.001), with a significantly decreased accuracy in the incongruent condition compared to the congruent condition (incongruent: mean+SEM=84%+0.70; congruent: mean+SEM=85%+0.60) and an increased accuracy on the approaching condition compared to avoiding condition (approach: mean+SEM=98.5%+0.20; avoid: mean+SEM=70.5%+1.10, Fig.1B). No significant interaction effect between congruence*action*group was found (F_(1,225)_=0.001, p=0.970).

For the response reaction time on the correct approach trials (e.g. Go trials), the mixed ANOVA analysis showed a significant interaction effect between group and congruence (F_(1,225)_=4.213, p=0.041), with the further Post-hoc tests showing that the reaction time in the incongruent condition was significantly higher than congruent condition for non-anxious group (incongruent: mean+SEM=322.127ms+5.856; congruent: mean+SEM= 314.456ms+5.604; p<0.001) but not the high-anxious group (incongruent: mean+SEM=320.988ms+5.196; congruent: mean+SEM=318.022ms+4.973; p=0.053, Fig.S1). This might indicate an over-generalized response when approaching positive stimuli and negative stimuli for the high-anxious group. In addition, in line with the findings reported by Bramson et al., (2023), the result also found a significant main effect of congruence (F_(1,225)_=21.530, p<0.001), with the reaction time in the incongruent condition was significantly higher than congruent condition (incongruent: mean+SEM=321.557ms+3.914; congruent: mean+SEM= 316.239ms+3.746).

3.2 BOLD level results

The neural congruency-effect across groups showed a significant recruitment of left DLPFC (SVC, Z=3.79, p_FWE_=0.015, voxels=40, x/y/z: -21, 24, 33) rather than FPl for the neural congruency-effect across groups (contrast: (HNG+ FG) > (HG+ FNG)). In addition, the results showed a significant activation in left MOG (middle occipital gyrus, MOG) extending to angular and SPG (superior parietal gyrus, SPG) on the whole brain level (initial thresholding p<0.001 uncorrected, cluster level, p_FWE_<0.05, see Table. S4).

Context-dependent activation analyses showed that both high-anxious and non-anxious groups showed similar activation patterns in FPl and DLPFC after SVC in different emotional contexts (see Table S1) and widespread brain activation in regions located at fronto-parietal, occipital and temporal lopes on the whole brain level (initial thresholding: p<0.001 uncorrected; cluster level, p_FWE_<0.05, see Fig. S2). In addition, no significant between-group differences were found in neither the negative (contrast: FG>FNG) nor positive context (contrasts: HNG>HG).

3.3 Functional connectivity results

Functional connectivity analyses were performed in the positive and negative context respectively with sgACC seeds defined from either positive or negative emotional action control (contrasts: HNG>HG; FG>FNG). For the sgACC seed defined from the positive contrast (HNG>HG), the results showed a positive connectivity between right sgACC and bilateral DLPFC but not FPl after SVC especially in the negative condition for high-anxious group (SVC: left DLPFC, Z=3.84, p_FWE_ =0.025, voxels=15, x/y/z: -24, 42, 27; right DLPFC, Z=4.06, p_FWE_ =0.012, voxels=23, x/y/z: 30, 42, 30). On the whole brain level, we found a positive connectivity between right sgACC and right ITG for high-anxious group in the negative context (Fig.S3A, details see Table. S5). In addition, while in the positive context there was a positive connectivity between right sgACC with ipsilateral caudal regions and left IFG and a negative connectivity with bilateral precuneus and left postcentral cortex for high-anxious group, the results showed a positive connectivity between right sgACC and fusiform, left MFG and a negative connectivity with bilateral precuneus for non-anxious group (Fig.S3B, details see Table. S5).

For the sgACC seed from the negative contrast (FG>FNG), the results showed a significant positive connectivity between left sgACC and bilateral DLPFC as well as a significant negative connectivity with bilateral FPl for the high-anxious group rather than the non-anxious group especially in the negative context after SVC. On the whole brain level while the results found a significant positive sgACC connectivity with left SFG, mOFC, MTG, MOG, right precuneus and angular regions and negative connectivity with bilateral mOFC and right cerebellum for high-anxious group, there is a negative connectivity between sgACC and right cerebellum for non-anxious group in the negative context (cluster level, p_FWE_<0.05, details see Table. S2).

3.4 Brain and scales association results

Functional connectivity strength (between sgACC and bilateral FPI and DLPFC) and scales association analyses including LSAS and total STAI scores were performed with age and gender as covariates respectively. After the correction for multiple comparisons (p<0.05/8=0.006), the results showed significant positive correlations between LSAS scores with sgACC-bilateral DLPFC connectivity strength in the negative context (sgACC-left DLPFC: r=0.193, p=0.002; sgACC-right DLPFC: r=0.177, p=0.004) but not with the sgACC-bilateral FPI connectivity strength (sgACC-left FPl: r=0.055, p=0.206; sgACC-right FPl: r=0.119, p=0.037). In addition, no significant correlations between STAI scores and the sgACC-FPI and sgACC-DLPFC connectivity were found (ps≥0.016).

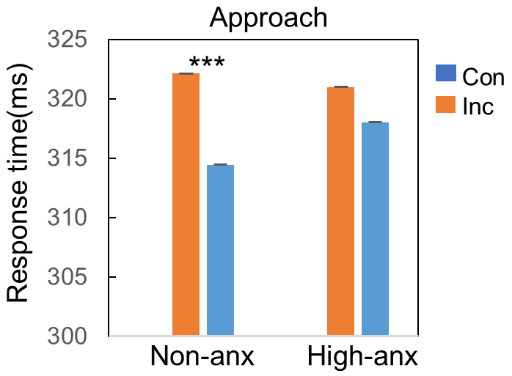

**Figure legend**

Fig. S1 The congruent effect on the response time for the correct approach trials (e.g. Go trials). The mixed ANOVA analysis showed a significant interaction effect between group and congruence (F_(1,225)_=4.213, p=0.041), with the further Post-hoc tests showing that the reaction time in the incongruent condition was significantly higher than congruent condition for non-anxious group (incongruent: mean+SEM=322.127ms+5.856; congruent: mean+SEM=314.456ms+5.604; p<0.001) but not the high-anxious group (incongruent: mean+SEM=320.988ms+5.196; congruent: mean+SEM=318.022ms+4.973; p=0.053). In addition, the result also found a significant main effect of congruence (F_(1,225)_=21.530, p<0.001), with the reaction time in the incongruent condition was significantly increased than congruent condition (incongruent: mean+SEM=321.557ms+3.914; congruent: mean+SEM=316.239ms+3.746). Note: Con: congruent; High-anx: high-social anxious group; Inc: incongruent; Non-anx: Non-social anxious group.*** means p<0.001.

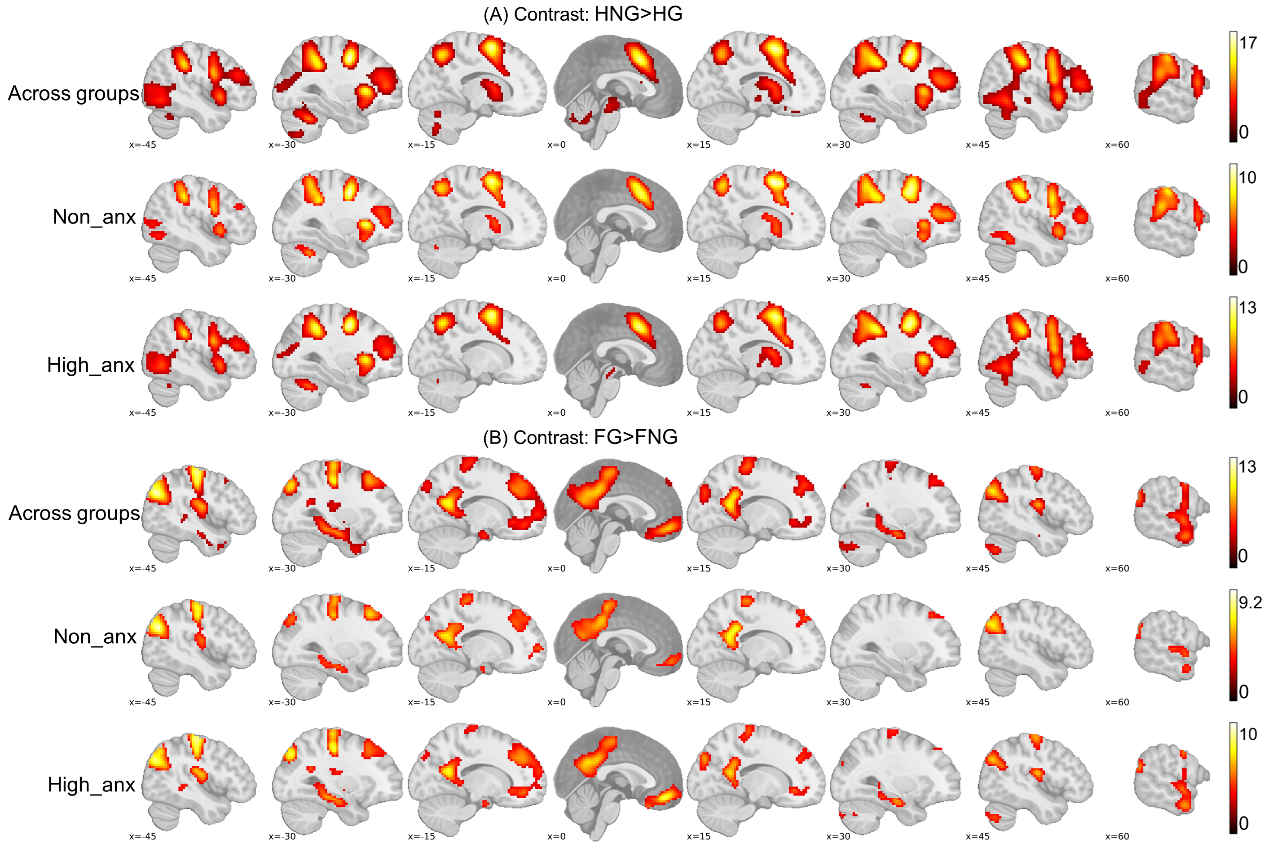

Fig. S2 Context-dependent activation mapping during action control on the whole brain level. (A) Brain activation mapping of action control in the positive context (contrast: HNG>HG) across groups as well as for both high-anxious and non-anxious group. (B) Brain activation mapping of action control in the negative context (contrast: FG>FNG) across groups as well as for both high-anxious and non-anxious group. All clusters passed the initial thresholding p<0.001 uncorrected, then cluster level p_FWE_ < 0.05. FWE, Family Wise Error. Note: High-anx: high-social anxious group; Non-anx: Non-social anxious group.

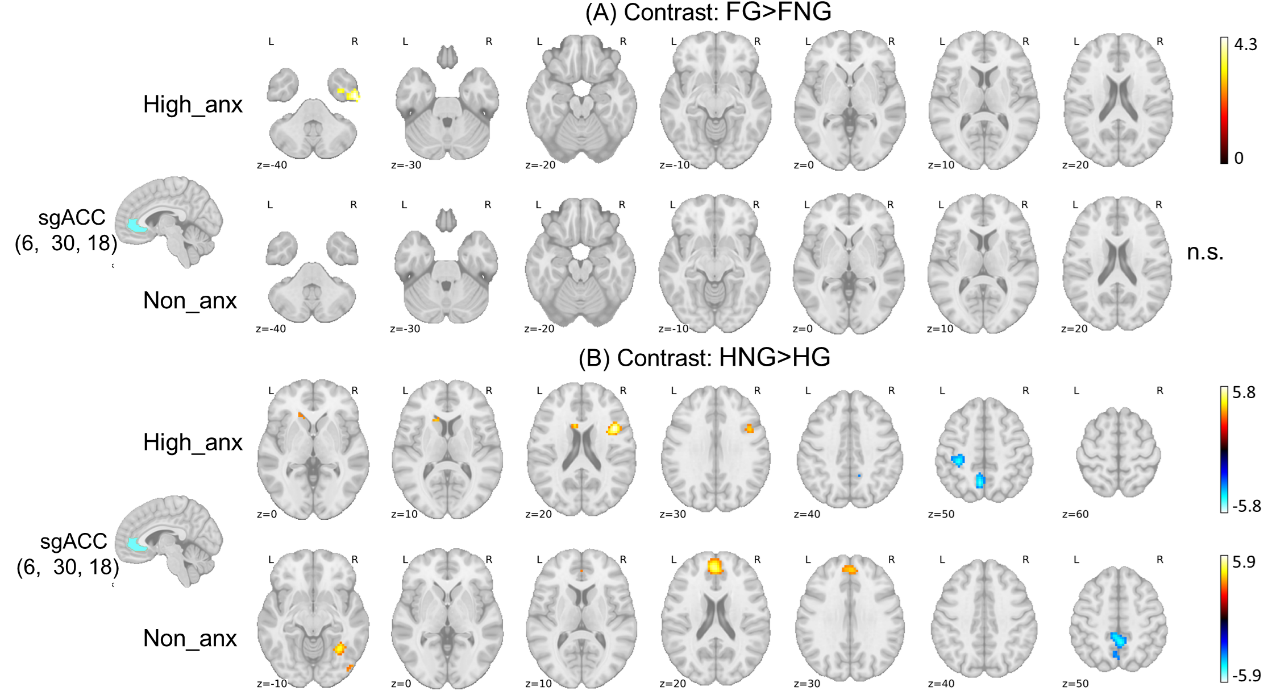

Fig. S3 Functional connectivity results for sgACC (seed defined from the contrast: HNG>HG) in the negative and positive context. (A)The results showed a positive connectivity between right sgACC and bilateral DLPFC especially in the negative condition for high-anxious group after SVC (SVC: left DLPFC, Z=3.84, p_FWE_ =0.025, voxels=15, x/y/z: -24, 42, 27; right DLPFC, Z=4.06, p_FWE_ =0.012, voxels=23, x/y/z: 30, 42, 30). On the whole brain level, the results showed a positive connectivity between right sgACC and right ITG for high-anxious group in the negative context. (B) While in the positive context there was positive connectivity between right sgACC with ipsilateral caudal regions and left IFG and a negative connectivity with bilateral precuneus and left postcentral cortex for high-anxious group, the results showed a positive connectivity between right sgACC and fusiform, left MFG and a negative connectivity with bilateral precuneus for non-anxious group. Note: High-anx: high-social anxious group; Non-anx: Non-social anxious group.

**Tables**

**Table S1. FPl and DLPFC activation in each group during behavioral control in either positive or negative context (contrasts: HNG>HG; FG>FNG).**

| Regions | Cluster K | Coordinates | | | t value |
| --- | --- | --- | --- | --- | --- |
|  |  | X | Y | Z |  |
| High-anxious group |  |  |  |  |  |
| Contrast: HNG>HG |  |  |  |  |  |
| l FPl | 140 | -30 | 48 | 12 | 6.37 |
| r FPl | 170 | 33 | 45 | 15 | 6.16 |
| l DLPFC | 57 | -30 | 39 | 27 | 5.18 |
| r DLPFC | 75 | 36 | 36 | 27 | 7.14 |
| Contrast: FG>FNG |  |  |  |  |  |
| l FPl | 32 | -21 | 57 | 0 | 4.19 |
| l DLPFC | 260 | -21 | 27 | 39 | 7.30 |
| r DLPFC | 94 | 18 | 39 | 36 | 4.80 |
| Non-anxious group |  |  |  |  |  |
| Contrast: HNG>HG |  |  |  |  |  |
| l FPl | 112 | -33 | 45 | 12 | 4.95 |
| r FPl | 150 | 27 | 48 | 15 | 6.37 |
| l DLPFC | 36 | -30 | 36 | 21 | 4.62 |
| r DLPFC | 54 | 33 | 36 | 24 | 6.12 |
| Contrast: FG>FNG |  |  |  |  |  |
| l FPl | 18 | -18 | 63 | 3 | 4.33 |
| l DLPFC | 224 | -24 | 27 | 48 | 6.58 |
| r DLPFC | 110 | 21 | 42 | 42 | 5.49 |

Note: Threshold: p<0.001 uncorrected, then small volume correction using the 15mm spheres as masks which is centered at the coordinates reported by Bramson et al., (2023), peak level p_FWE_ < 0.05. DLPFC, dorsolateral prefrontal cortex; FPl, lateral frontopolar cortex; FWE, Family Wise Error; l, left; r, right.

**Table S2. Left sgACC (seed defined from contrast: FG>FNG) connectivity results for high-anxious and non-anxious group in the negative context on the whole brain level.**

| Regions | Cluster K | Coordinates | | | t value |
| --- | --- | --- | --- | --- | --- |
|  |  | X | Y | Z |  |
| Results for high-anxious group | | | | | |
| Positive connectivity with sgACC | | | | | |
| l SFG | 283 | -21 | 39 | 39 | 5.53 |
|  |  | -15 | 48 | 39 | 5.38 |
|  |  | -21 | 27 | 45 | 4.62 |
| r mOFC | 139 | 0 | 51 | 6 | 3.99 |
|  |  | 0 | 51 | -15 | 3.98 |
|  |  | -3 | 60 | 3 | 3.82 |
| r Precuneus | 208 | 6 | -54 | 15 | 4.20 |
|  |  | 0 | -60 | 30 | 4.03 |
|  |  | -9 | -51 | 30 | 3.91 |
| l MOG | 408 | -39 | 0 | 3 | 5.46 |
|  |  | -57 | -18 | 33 | 5.29 |
|  |  | -63 | -18 | 21 | 5.21 |
| r Angular | 101 | 48 | -75 | 33 | 5.89 |
|  |  | 57 | -63 | 15 | 4.46 |
| l MTG | 103 | -60 | -33 | -6 | 4.36 |
|  |  | -48 | -48 | -6 | 4.32 |
| Negative connectivity with sgACC | | | | | |
| l mOFC | 288 | 18 | 27 | -21 | 6.07 |
|  |  | 21 | 54 | -9 | 4.83 |
|  |  | 9 | 24 | -12 | 4.69 |
| r mOFC | 265 | -21 | 39 | -21 | 5.31 |
|  |  | -18 | 33 | -9 | 5.17 |
|  |  | -24 | 42 | -6 | 4.85 |
| r Cerebellum | 115 | 33 | -51 | -45 | 5.23 |
|  |  | 24 | -42 | -54 | 4.58 |
| Results for non-anxious group | | | | | |
| Negative connectivity with sgACC | | | | | |
| r Cerebellum | 96 | 15 | -78 | -45 | 5.01 |

Note: All clusters passed the initial thresholding p<0.001 uncorrected, then cluster level p_FWE_ < 0.05. FWE, Family Wise Error; l, left; mOFC, medial orbital frontal cortex; MOG, Middle occipital gyrus; MTG, Middle temporal gyrus; r, right; SFG, Superior frontal gyrus; sgACC, subgenual Anterior Cingulate Cortex.

**Table S3. sgACC activation across groups during action control in either positive or negative contexts (contrasts: HNG>HG; FG>FNG).**

| Regions | Cluster K | Coordinates | | | t value |
| --- | --- | --- | --- | --- | --- |
|  |  | X | Y | Z |  |
| contrast: HNG>HG |  |  |  |  |  |
| r sgACC | 10 | 6 | 30 | 18 | 6.67 |
|  |  | 9 | 36 | 15 | 4.98 |
| contrast: FG>FNG |  |  |  |  |  |
| l sgACC | 35 | -3 | 45 | -9 | 4.93 |
|  |  | -3 | 33 | -15 | 4.68 |
|  |  | -6 | -39 | -12 | 4.81 |

Note: Threshold: initial thresholding p<0.001 uncorrected, then small volume correction with the sgACC mask from the Human brainnetome atlas (Fan et al., 2016), peak level p_FWE_ < 0.05. FWE, Family Wise Error; l, left; r, right; sgACC, subgenual anterior cingulate cortex.

**Table S4. The neural congruency-effect across groups on the whole brain level.**

| Regions | Cluster K | Coordinates | | | t value |
| --- | --- | --- | --- | --- | --- |
|  |  | X | Y | Z |  |
| l MOG extending to angular and SPG | 247 | -36 | -78 | 36 | 4.54 |
|  |  | -39 | -60 | 30 | 3.62 |
|  |  | -18 | -75 | 45 | 3.31 |

Note: All clusters passed the thresholding at: initial thresholding p<0.001 uncorrected, cluster level p_FWE_ < 0.05. FWE, Family Wise Error; l, left; MOG, middle occipital gyrus; SPG, superior parietal gyrus.

**Table S5. Right sgACC (seed defined from contrast: HNG>HG) connectivity results for high-anxious and non-anxious group on the whole brain level.**

| Regions | Cluster K | Coordinates | | | t value |
| --- | --- | --- | --- | --- | --- |
|  |  | X | Y | Z |  |
| Results for high-anxious group in the negative context | | | | | |
| Positive connectivity with sgACC | | | | | |
| r ITG | 108 | 54 | -12 | -39 | 4.34 |
|  |  | 39 | -3 | -42 | 4.28 |
|  |  | 48 | -6 | -48 | 3.89 |
| Results for high-anxious group in the positive context | | | | | |
| Positive connectivity with sgACC | | | | | |
| r IFG | 132 | 42 | 9 | 24 | 5.77 |
|  |  | 45 | 30 | 15 | 3.99 |
| l Caudate | 89 | -12 | 18 | 15 | 4.75 |
|  |  | -30 | 27 | 15 | 4.13 |
|  |  | -24 | 15 | 15 | 3.53 |
| Negative connectivity with sgACC | | | | | |
| l Precuneus | 107 | -6 | -60 | 51 | 4.74 |
|  |  | 9 | -54 | 45 | 4.13 |
| l Postcentral cortex | 85 | -33 | -33 | 51 | 4.50 |
| Results for non-anxious group in the positive context | | | | | |
| Positive connectivity with sgACC | | | | | |
| l MFG | 217 | -3 | 51 | 21 | 5.20 |
| r Fusiform | 125 | 33 | -57 | -6 | 4.83 |
|  |  | 48 | -78 | -6 | 4.14 |
| Negative connectivity with sgACC | | | | | |
| l Precuneus | 336 | 0 | -48 | 57 | 5.95 |
|  |  | -3 | -57 | 60 | 5.55 |

Note: All clusters passed the initial thresholding p<0.001 uncorrected, then cluster level p_FWE_ < 0.05. FWE, Family Wise Error; IFG, inferior frontal gyrus; ITG, inferior temporal gyrus; l, left; MFG, middle frontal gyrus; r, right; sgACC, subgenual Anterior Cingulate Cortex.
